# Application of metagenomic sequencing to improve sessile serrated lesion detection using the faecal immunochemical test

**DOI:** 10.64898/2026.11.04.764411

**Authors:** Bradley J. Hart, Geraldine Laven-Law, Clarice M. Harker, Michael P. Doane, Rachel Parker, Martha Menberu, Charles Cock, Robert A. Edwards, Erin L. Symonds, Jean M. Winter, Michael J. Roach

## Abstract

Most population colorectal cancer (CRC) screening programs use the faecal immunochemical test (FIT), which quantifies occult haemoglobin in faeces and has proven to be effective in detecting and reducing CRC incidence and mortality. Sessile serrated lesions (SSLs) account for up to 30% of sporadic CRC, yet FIT has low sensitivity to detect them. Faecal microbial biomarkers may improve their detection, but FIT samples have not previously been used to study the SSL microbiome. We asked whether FIT samples preserve the faecal microbiome as accurately as fresh collection, and whether they can reveal a microbial signature of SSL. We first reanalysed a published dataset of paired FIT and fresh faecal samples from the same participants. Donor identity explained 63.5% of compositional variance compared with 16.7% for collection method, indicating that FIT preserves the donor-specific signature on which biomarker discovery depends. We then generated shotgun metagenomes from FIT samples of 24 adults scheduled for diagnostic colonoscopy, twelve with at least one clinically significant SSL and twelve with no neoplasia (NN) and compared each group with a published fresh faeces cohort matched on colonoscopy outcome. Collection method explained 6.3% to 7.6% of variance, and no individual species was significantly differentially abundant between methods. Between SSL and NN FIT samples, community structure did not significantly differ, and conservative differential abundance analysis returned no results. More permissive count-based models suggested a depletion of fermentative commensals in SSL. Functionally, SSL samples encoded significantly fewer KEGG orthologs and less polyamine biosynthesis capacity, while orthologs carried by several *Bifidobacterium* species were more abundant. Many depleted functions reflected *Enterobacteriaceae* gene content concentrated in a few control samples. No single feature was diagnostic on its own. A multi-feature signature, confirmed in larger cohorts and applied to samples that screening programs already collect, may increase the detection rate of SSLs.

## INTRODUCTION

Colorectal cancer (CRC) is the third most diagnosed cancer and the second leading cause of cancer-related death worldwide (1). Most CRCs develop slowly, over approximately 10–15 years, from precancerous lesions such as adenomas and sessile serrated lesions (SSLs), so CRC is largely preventable if these lesions are detected and removed at colonoscopy. The faecal immunochemical test (FIT) is used worldwide as a CRC screening tool and effectively reduces CRC-related mortality (2). It detects occult blood in faeces using antibodies specific to haemoglobin with a sensitivity for CRC at 52–94% (3). FIT is considerately less sensitive for SSLs. These are flat lesions with a characteristic saw-tooth histological appearance and are thought to give rise to up to 30% of all CRCs (4), yet FIT only detects up to 12.3% of them (5). In part, this reflects their biology. SSLs are hypo-vascular and bleed little, and their predominantly right-sided location means that any blood they shed is degraded during colonic transit. Where FIT does identify an SSL, the detection is thought to be largely incidental, arising from a synchronous adenoma or a benign bleeding source such as haemorrhoids rather than from the SSL itself (5–7). Therefore, a different type of biomarker is required.

The gut microbiome offers a complementary source of screening information. Castellarin et al. (8) used high-throughput sequencing to show that colorectal tumours were enriched for *Fusobacterium nucleatum*, and Zackular et al. (9) showed that faecal microbial profiles could distinguish healthy individuals from those with adenomas or carcinomas, proposing the microbiome as a screening tool. Far less is known about SSLs. The studies that have included them were small, mostly utilising 16S rRNA metabarcoding or mucosal samples, and reported differing taxa (10–12), while the functional capacity of the community received little attention. Repurposing the FIT sample would allow microbial markers to be measured in specimens that screening programs already collect. However, the FIT buffer is designed to stabilise haemoglobin rather than microbial DNA, and samples can spend up to two weeks at ambient temperature before reaching the laboratory, so the device must first be shown not to distort the community. We therefore first reanalysed a published shotgun dataset in which FIT and fresh faecal samples were collected from the same participants (13), measuring the collection-method effect within each person. We then compared shotgun metagenomes from our own FIT samples with two published fresh faeces cohorts, matched on colonoscopy findings so that collection method was not confounded with lesion status. Finally, we compared FIT samples from participants with a clinically significant SSL to those with no neoplasia (NN) to identify taxonomic and functional differences associated with SSL.

## MATERIALS AND METHODS

### Study population

This analysis utilised samples from a subset of participants in a larger prospective study of adults (≥ 18 years) scheduled for diagnostic colonoscopy within the Southern Adelaide Local Health Network (ACTRN12620000668909). Invitations were mailed with an information sheet, a consent form and a home collection kit containing two FIT sampling tubes (OC-Auto Sampling Bottle 3, EIKEN CHEMICAL CO., LTD. Japan), collection sheets, test completion instructions, and a reply-paid envelope. Diet and medication were unrestricted. Demographic information was collected from participants via health questionnaires and hospital records. Ethical approval was obtained from the Southern Adelaide Clinical Human Research Ethics Committee (#35.20). The trial was prospectively registered with the Australian and New Zealand Clinical Trials Registry (Registry ID #ACTRN12620000668909). The study followed the Declaration of Helsinki and all participants provided written and informed consent.

### Faeces sample collection

Each OC-Sensor FIT device obtains approximately 10 mg of faeces sampled from the outer surface of the faeces and contains 2 mL of haemoglobin stabilisation buffer. Participants sampled from two separate faecal samples within 90 days prior to colonoscopy, recording sample collection time and date. Completed tests were stored in a cool place prior to mail return and then mailed to the Flinders Centre for Innovation in Cancer laboratory, South Australia, Australia.

### FIT haemoglobin analysis

De-identified FIT devices were stored at 4°C upon arrival. FIT samples were analysed for haemoglobin using the automated OC-Sensor DIANA or OC-Sensor PLEDIA faecal testing devices (EIKEN CHEMICAL CO., LTD.) as per the manufacturer’s protocols, within 14 days of the first faecal sample collection. Samples were deemed positive for haemoglobin if either of the two completed FIT devices contained ≥20 µg haemoglobin/g faeces, consistent with the Australian National Bowel Cancer Screening Program (14). FIT results were blinded to participants and clinicians. Following haemoglobin analysis, FITs were wrapped in parafilm and stored at −20°C until further use.

### Colonoscopy

Colonoscopies were conducted or supervised by nationally accredited gastroenterologists or colorectal surgeons at hospitals affiliated with the Southern Adelaide Local Health Network. Only colonoscopies consistent with quality definitions in the Australian Colonoscopy Clinical Care Standard (15) were included, with procedures including complete intubation (e.g. to the caecum, terminal ileum, or appendiceal orifice), and with sufficient bowel preparation with a Boston Bowel Preparation Score (16) of 2 or more in each colonic segment. Polyps found at colonoscopy were retrieved using standard techniques for histopathological analysis, with histopathology reported by hospital pathologists.

Participants were classified by the most advanced lesion found. The no neoplasia (NN) group had normal findings or benign lesions such as haemorrhoids. The SSL group had at least one SSL ≥10 mm, or an SSL of any size with cytological dysplasia (Supplementary Table S1).

Participants with an SSL meeting this definition were assigned to the SSL group regardless of synchronous conventional adenomas. The terminology related to CRC has been redefined over time (17), so to avoid confusion with other similar terms to SSL, we provide definitions in Supplementary Table S1.

### DNA extraction, library preparation, and sequencing

Frozen FIT devices were thawed on ice, opened using shears, and 250 µL of the contents (faecal matter in buffer) was homogenised (Precellys Evolution, Thermo Fisher) and extracted with the ZymoBIOMICS DNA Miniprep Kit (Zymo Research). DNA was quantified by Qubit (Thermo Fisher). Extracts from each participant’s two tubes were pooled, and each sample of 15 ng pooled DNA was submitted to the Australian Genome Research Facility for library preparation with the DNA Preparation M kit (Illumina) and subsequent whole-genome metagenomics sequencing on the NextSeq 1000/2000 (Illumina) with both 300PE and 150PE chemistry.

### Sequence preprocessing and profiling

Both 150PE and 300PE sequence results were concatenated, and all samples were processed through an identical pipeline. Reads were quality controlled with Trimnami v0.1.3 (18), which wraps Fastp v0.23.2 (19) for adapter and quality trimming while also discarding pairs mapping to the human genome. Statistics related to post-processed sequences are available in Supplementary Table S2.

Species-level composition was assigned from the cleaned reads with MetaPhlAn4 v4.2.4 (20) which maps clade-specific marker genes and reports relative abundances using its species-level genome bin (SGB) database with NCBI-based taxonomy. Merged tables were parsed to the seven standard ranks and filtered to species-level clades, defined as rows with an s assignment and no strain-level t prefix. Functional potential was profiled with HUMAnN3 v3.9 (21) against the default ChocoPhlAn and UniRef90 databases, giving MetaCyc pathway and KEGG orthologue (KO) tables in both copies-per-million (CPM) and reads-per-kilobase (RPK) normalisations, with unmapped and unintegrated features removed. Because GTDB reorganises several genera relative to National Center for Biotechnology Information (NCBI) classifications, comparisons with earlier literature account for reclassifications such as Bacteroides to *Phocaeicola* and Prevotella to *Segatella*.

### Reference cohorts and study comparisons

To assess whether FIT-collected samples yield microbial profiles comparable to those obtained from directly collected stool, and are therefore suitable for downstream comparative analysis, three comparisons were performed: one using paired samples from the same individuals and two against independent reference cohorts. The first reanalysed publicly available data from Byrd et al. (13), in which paired FIT and fresh faecal samples were collected from the same participants, so that the effect of collection method could be measured within each person and our bioinformatics process could be validated against a published result. The second and third compared the 24 FIT samples from the current study with 24 fresh faecal samples from two published cohorts (Table 1), split by neoplasia status so that collection method was not confounded with lesion status: 12 NN FIT samples against 12 neoplasia-free fresh samples from Piccinno et al. (22), and 12 SSL FIT samples against 12 SSL fresh samples from Lee et al. (23). Neoplasia status was confirmed at colonoscopy in both cross-cohort comparisons. Byrd et al. recruited healthy volunteers. All external datasets were processed through the same pipeline as our FIT samples.

**Table 1.** Participant demographics and sample characteristics across cohorts included in cross-cohort comparisons. FIT, faecal immunochemical test; SSL, sessile serrated lesion; IQR, interquartile range; N.A., not assessed; n.d., no data, SD, standard deviation; Hb, haemoglobin; ᵃ Lee et al. (2023) reported age as decade-wide bins. Statistics are approximated from bin midpoints; exact continuous values were not available from the Sequence Read Archive metadata. ᵇ Lee et al. (2023) Phenotype field ‘SSL only’ denotes sessile serrated lesions as the sole lesion type; all 12 participants had confirmed SSL at colonoscopy.

| Characteristic | FIT Cohort |  | Comparison benign cohort | Comparison SSL cohort |
|---|---|---|---|---|
|  | SSL | No neoplasia | Piccinno et al. (2025) | Lee et al. (2023) |
|  | FIT (n = 12) | FIT (n = 12) | Benign faeces (n = 12) | SSL faeces (n = 12) |
| Study design |  |  |  |  |
| Source publication | FIT Cohort |  | Piccinno et al. (2025) | Lee et al. (2023) |
| Country | Australia |  | Czechia | Singapore |
| Sample type | FIT (thawed) |  | Fresh (faeces) | Fresh (faeces) |
| Sequencing method | Shotgun metagenomics |  | Shotgun metagenomics | Shotgun metagenomics |
| Instrument | Illumina NextSeq |  | Illumina | Illumina HiSeq 2500 |
| Accession | PRJNA1522997 |  | PRJEB72525 | PRJNA784939 |
| Demographics |  |  |  |  |
| Total participants, n | 12 | 12 | 12 | 12 |
| Male, n (%) | 3 (25.0%) | 6 (50.0%) | 6 (50.0%) | 6 (50.0%) |
| Female, n (%) | 9 (75.0%) | 6 (50.0%) | 6 (50.0%) | 6 (50.0%) |
| Age (years) |  |  |  |  |
| Mean ± SD | 55.9 ± 15.3 | 59.1 ± 15.7 | 57.4 ± 14.4 | ~57.8 (midpoint est.)<sup>a</sup> |
| Median (IQR) | 57.0 (48.0–64.2) | 61.0 (48.0–69.8) | 61.5 (51.0–69.0) | 55–64 (bin range)<sup>a</sup> |
| Range | 25–85 | 26–83 | 23–74 | 20–79 (bin range)<sup>a</sup> |
| Body mass index (kg/m²) |  |  |  |  |
| Median (IQR) | 29.9 (27.5–33.8) | 26.1 (23.1–27.1) | 27.9 (23.4–33.5) | n.d. |
| Lesion characteristics |  |  |  |  |
| SSL, n (%) | 12 (100%) | 0 (0%) | 0 (0%) | 12 (100%)<sup>b</sup> |
| Lesion size ≥10 mm, n (%) | 12 (100%) | 0 (0%) | 0 (0%) | n.d. |
| Cytological dysplasia, n (%) | 2 (16.7%) | 0 (0%) | 0 (0%) | n.d. |
| Synchronous advanced adenoma, n (%) | 5 (41.7%) | 0 (0%) | 0 (0%) | n.d. |
| FIT metrics |  |  |  |  |
| FIT positive, n (%) | 4 (33.3%) | 3 (25.0%) | N.A. | N.A. |
| FIT Hb — median (IQR) | 4.8 (3.9–42.3) | 0.4 (0.2–12.3) | N.A. | N.A. |
| Sample age — median (IQR) | 9.5 (7.0–12.0) | 9.0 (6.8–11.0) | 0 | 0 |

### Filtering, Alpha and beta diversity, and ordination

For the MetaPhlAn4 analyses a prevalence filter was applied before normalisation, retaining taxa present at non-zero abundance in more than 10% of samples. For the HUMAnN3 analyses, a feature had to reach at least 25% prevalence in either group, computed on the RPK layer, after which features with constant detected values within either group were removed. Where count-based models were used, relative abundances were converted to pseudo-counts by total-sum scaling to proportions and multiplying by 10⁶ before rounding. Within-sample diversity used Shannon and Simpson indices for taxonomy, extended to observed richness, inverse Simpson and Pielou’s evenness for function. Groups were compared by Wilcoxon rank-sum test with Cliff’s delta as effect size, absolute values below 0.147 were treated as negligible and below 0.33 as small (24). Taxonomic between-sample structure was measured with Bray-Curtis dissimilarity on relative abundances and with Aitchison distance (the Euclidean distance between centred log-ratio (CLR) transformed abundances), both ordinated by principal coordinates analysis. Functional profiles were ordinated by non-metric multidimensional scaling of Bray-Curtis dissimilarity on pathway CPM and by principal components analysis of CLR-transformed pathway RPK. Group separation was tested by PERMANOVA (adonis2, 999 permutations), which asks whether the average community differs between groups, with PERMDISP used to ask whether within-group variability differs instead. For the Byrd et al. (13) data, variance was partitioned by PERMANOVA with sequential sums of squares fitting participant before method, and alpha diversity by two-way ANOVA with participant and method as main effects.

### Differential abundance

Species-level differential abundance was tested with ALDEx2 v1.42.0 (25) in all four comparisons. ALDEx2 accounts for the compositional nature of sequencing data by applying a centred log-ratio transformation to 128 Monte Carlo Dirichlet instances, here with all features as the reference set, and significance was set at a Benjamini-Hochberg adjusted p < 0.05. In the paired Byrd et al. (13) comparison, tests were paired so that each participant’s FIT sample was matched to their own fresh sample. In the two cross-cohort comparisons, NN FIT against NN fresh and SSL FIT against SSL fresh, a species was considered altered by collection method if it reached significance under either Welch’s t-test or the Wilcoxon rank-sum test. The fourth comparison, FIT SSL against FIT NN, addresses the question of the study and was assessed by the Wilcoxon rank-sum test alone.

As ALDEx2 is conservative at 12 samples per group, the studies’ FIT SSL and FIT NN profiles were also analysed with DESeq2 v1.50.2 (26) and edgeR v4.8.2 (27), negative binomial models that detect more features in microbiome benchmarking at the cost of a higher false positive rate (28). DESeq2 calls were made at an adjusted p < 0.05, and edgeR used TMM normalisation with a quasi-likelihood F-test at FDR < 0.05. Species called by both methods with a concordant direction of change were retained as an exploratory set describing directional trends below the ALDEx2 threshold, not as differentially abundant features.

HUMAnN3 pathways and KOs were tested with three compositional methods: LinDA (microbiomestat v1.4) (25) which fits a linear model to log-ratio transformed data, MaAsLin3 v1.2.0 (26), which models total-sum-scaled, log-transformed features and ANCOM-BC2 v2.12.1 (27), applied to the RPK layer with Benjamini-Hochberg adjustment. A false discovery rate threshold of 0.25 preserved sensitivity at 12 samples per group, and features called significant by two or more methods formed the consensus set. To identify the taxa carrying each functional difference, the HUMAnN3 stratified output was used to divide the difference in mean CPM between groups for every consensus feature into per-species contributions (Table S5). Breadth was summarised as the effective number of contributing species (the inverse Simpson index of the positive per-species shares), and a feature was considered broadly carried when it involved three or more effective species and no single species contributed half or more of the difference. Because many functional shifts were driven by *E. coli* in two NN samples, interpretation focuses on broadly carried features.

## RESULTS

### Study Population

In our selected subset, faecal haemoglobin measured by FIT did not discriminate SSL from benign colonoscopy outcomes. Of the twenty-four FIT samples, four of the twelve participants with an SSL (33%) were FIT-positive, one of whom also had an advanced adenoma, compared with three of twelve (25%) with a benign outcome (p = 0.65; Table 1). All twelve SSLs were ≥ 10 mm, two showed dysplasia, and five participants also had a synchronous advanced adenoma (Table 1).

### Reanalysis of FIT and fresh faecal collection in a paired dataset confirms consistent microbiome profiles

Reanalysis of the Byrd et al. (13) paired sequences found that FIT was comparable to fresh faeces across three within-sample diversity measures (Shannon index, Simpson index, and the number of observed species, Figure 1a). Per-participant Bray-Curtis similarity which compares whole communities between samples, ranged from 19.3% to 54.3% across the 12 participants pairs (Figure 1b). Inter-participant differences dominated the paired comparison. Donor identity accounted for 63.5% (Bray-Curtis) and 67.1% (Aitchison) of total variance, compared with 16.7% and 13.9% for collection method, with the residual explaining 19.9% and 19.0% (Figure 1c). The method effect was significant under both metrics (p = 0.001, 999 permutations) but was approximately a quarter of the inter-participant signal. The method effect was directional. In all 12 participants, the FIT sample carried more *Bacteroidota* and less *Firmicutes* than the frozen sample (median *Bacteroidota* 46% versus 3%; Figure 1e), and only two pairs reached 50% Bray-Curtis similarity (Figure 1b). Taxa detected by only one method were present at very low relative abundance (< 0.3%), where detection may reflect sampling chance (Figure 1d; Supplementary Table S3).

**Figure 1.**
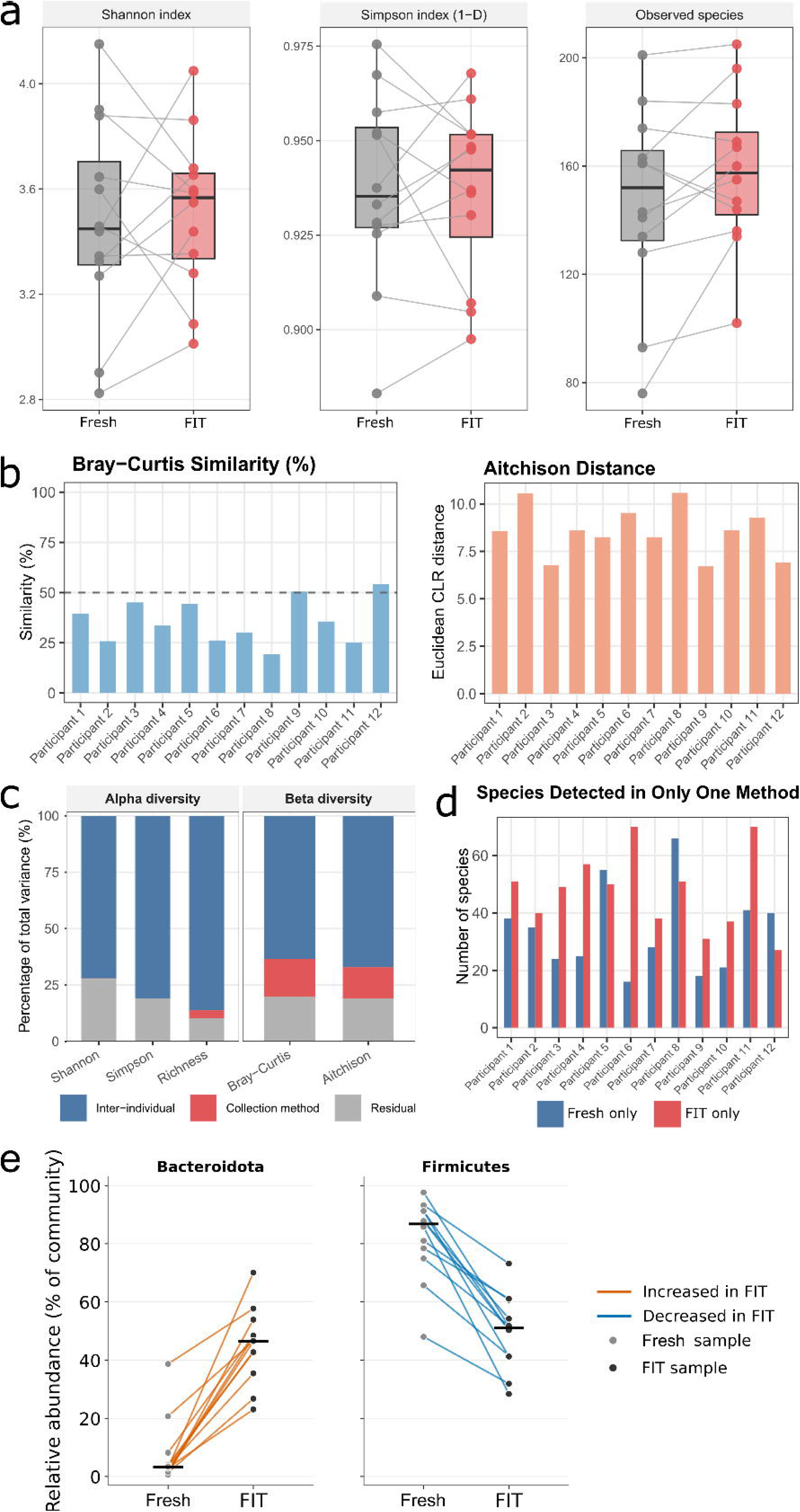
Collection method contributes far less to metagenomic variance than inter-individual differences. Published data consists of 12 paired FIT and fresh faeces samples (13). **a)** Within-sample alpha diversity by collection method, showing the Shannon index, Simpson index (1-D) and observed species richness. Each point is one sample and grey lines join the paired fresh and FIT samples from the same sample; fresh samples are shown in grey, FIT in red. **b)** Per-sample Bray-Curtis similarity with dashed line marking the 50% threshold (*left*), and Aitchison (Euclidean CLR) distance (*right*) between each participants two samples, quantifying the within-person divergence attributable to collection method. **c)** Variance partitioning for alpha diversity with balanced two-way ANOVA (*left*) and beta diversity with PERMANOVA on the full distance matrix (*right*). Both are normalised to 100% of total variance. Inter-individual variation accounts for 72.2% (Shannon), 81.0% (Simpson) and 89.4% (richness) of alpha-diversity variance and 63.5% (Bray-Curtis) and 67.1% (Aitchison) of beta-diversity variance, while collection method accounts for 3.6% or less of alpha-diversity variance and 16.7% or less of beta-diversity variance. **d)** Number of species detected by only one collection method per sample, coloured by the method of exclusive detection. **e)** Relative abundance in paired direct and FIT samples of Bacteroidota (*left*) and Firmicutes (*right*).

Alpha diversity was almost entirely determined by the donor, with collection method explaining 0.3% (Shannon), 0.0% (Simpson) and 3.6% (observed richness) of variance, against 72.2%, 81.0% and 89.4% for donor identity (Figure 1c).

### FIT samples consistent with published fresh faecal microbiomes

Our cohort FIT, and published fresh faecal samples were compared separately within the NN and SSL groups (Table 1). Alpha diversity did not differ significantly between FIT and fresh faeces in the NN comparison or in the SSL comparison (Table S6). Ordination of Bray-Curtis and Aitchison distances produced overlapping 95% confidence ellipses in both comparisons (Figure 2a; Figure 3a). Collection method reached significance under PERMANOVA in both cohorts and under both distances (NN: Bray-Curtis p = 0.009; Aitchison p = 0.014; SSL: Bray-Curtis p = 0.011; Aitchison p = 0.012), and explained 6.3% to 7.6% of total variance.

**Figure 2.**
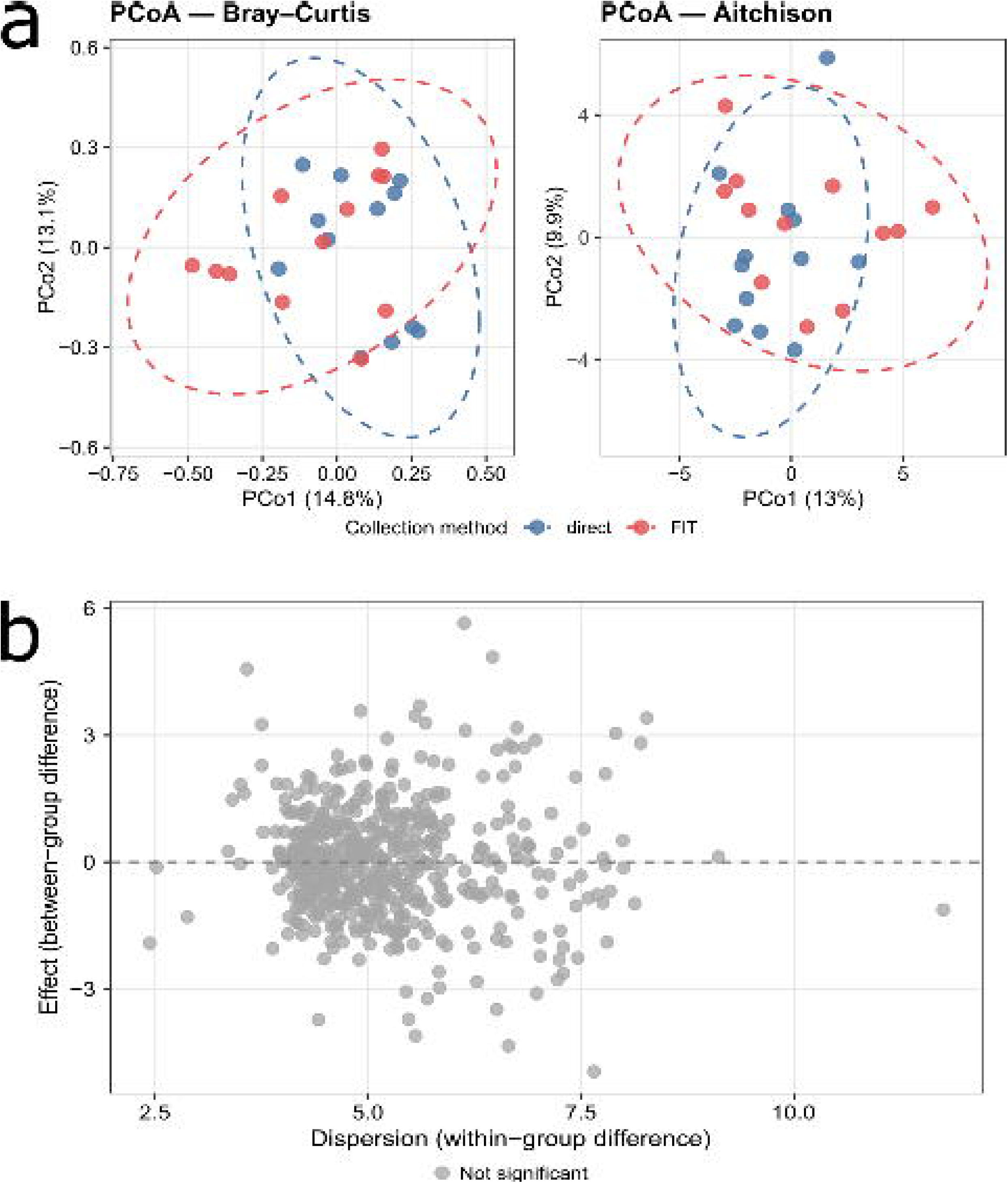
FIT and freshly collected faeces metagenomes are comparable in neoplasia-free donors. **a)** Principal coordinates analysis (PCoA) comparing 12 FIT-collected metagenomes (current study) with 12 fresh faeces metagenomes from a published neoplasia-free cohort (22), showing PCoA of Bray-Curtis dissimilarity computed on total sum scaling (TSS)-normalised species abundance profiles (*left*), and PCoA of Aitchison distance computed on centred log-ratio (CLR)-transformed species profiles (*right*). Ellipses denote 95% confidence intervals. **b)** ALDEx2 effect plot, showing between-group difference against within-group dispersion. Each point is one taxon; no taxon met the Benjamini-Hochberg corrected significance threshold (q < 0.05).

**Figure 3.**
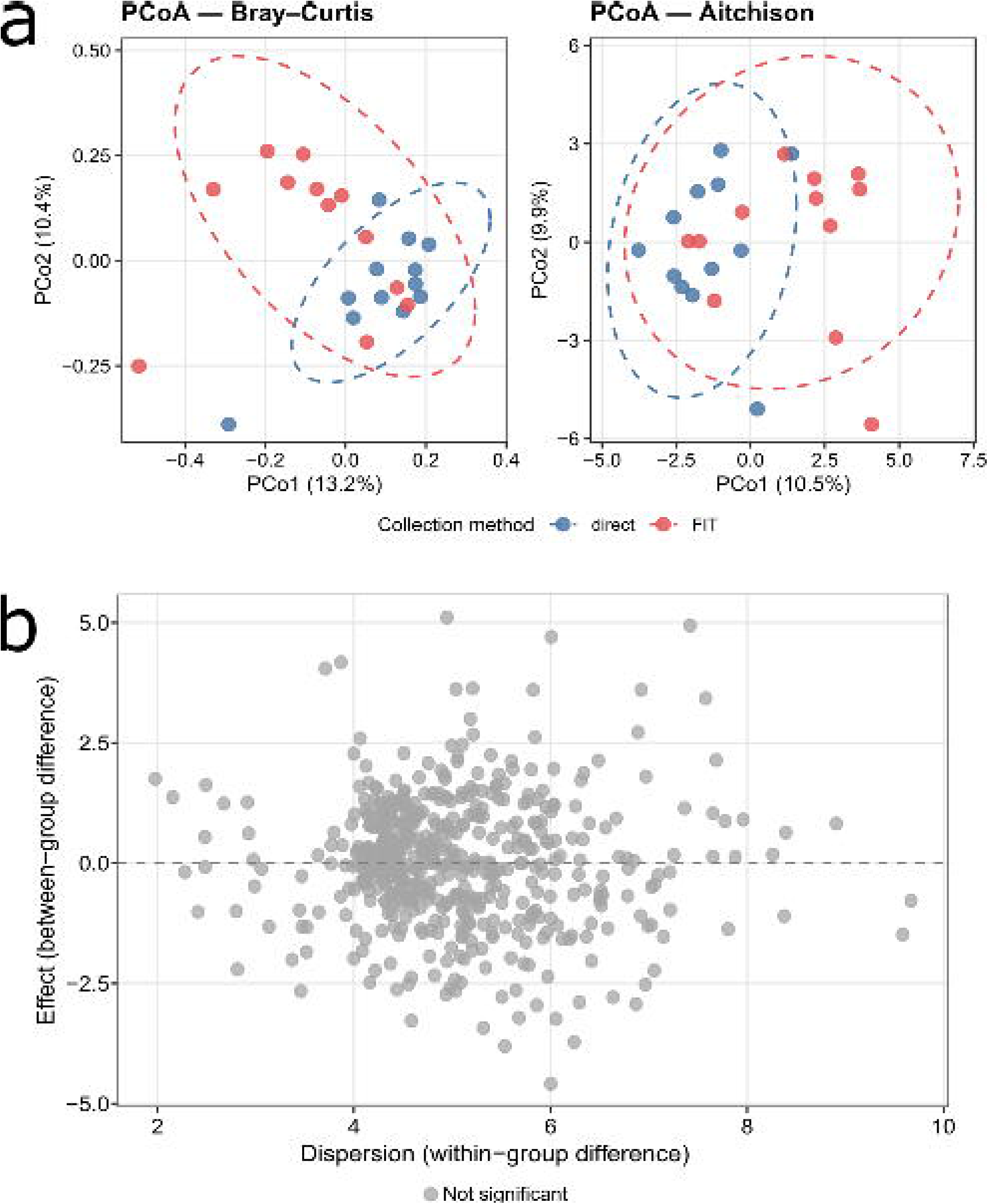
FIT and freshly collected faeces metagenomes are compositionally equivalent in donors with a sessile serrated lesion. **a)** Principal coordinates analysis (PCoA) comparing 12 FIT-collected metagenomes (current study) with 12 fresh faeces metagenomes from a published sessile serrated lesion cohort (23), showing PCoA of Bray-Curtis dissimilarity computed on total sum scaling (TSS)-normalised species abundance profiles (*left*), and PCoA of Aitchison distance computed on centred log-ratio (CLR)-transformed species profiles (*right*). Ellipses denote 95% confidence intervals. **b)** ALDEx2 effect plot, showing between-group difference against within-group dispersion. Each point is one taxon; no taxon met the Benjamini-Hochberg corrected significance threshold (q < 0.05).

Within-group dispersion was homogeneous throughout (PERMDISP NN: Bray-Curtis p = 0.451, Aitchison p = 0.724; SSL: Bray-Curtis p = 0.777, Aitchison p = 0.649), so the PERMANOVA result reflects a shift in centroid rather than a difference in spread. ALDEx2 identified no differentially abundant species in either comparison (Table S6; Figures 2b, 3b). Because each fresh cohort was sampled on another continent, this variance is shared between collection method and study of origin, so 6.3% to 7.6% is an upper bound on the FIT effect rather than a measurement of it. The two methods recovered closely overlapping species sets; Of the 534 species retained after prevalence filtering in the NN comparison, 467 were found by both methods (511 in FIT, 490 in fresh faeces), and of the 536 detected in the SSL comparison, 462 were shared (483 in FIT, 515 in fresh faeces), so neither method consistently contributed more taxa than the other. *Lachnospiraceae* was the most abundant family in FIT samples from both cohorts and in the SSL fresh samples, in NN fresh samples, *Bacteroidaceae* (21.7%) slightly exceeded *Lachnospiraceae* (18.9%) (Table 2). No individual species differed between collection methods. Some families differed in mean abundance between cohorts; the high *Bifidobacteriaceae* abundance in SSL FIT samples (11.9%) was not seen in NN FIT samples (2.6%), indicating a group difference rather than an effect of FIT collection.

**Table 2.** Five most abundant families per cohort, with mean relative abundance by collection method. Values are means of species-level abundances after prevalence filtering.

| Cohort | Family | Fresh faeces (mean %) | FIT (mean %) | Overall (mean %) |
|---|---|---|---|---|
| Non-neoplastic (Piccinno et al.) | *Lachnospiraceae* | 18.9 | 24.2 | 21.6 |
|  | *Bacteroidaceae* | 21.7 | 13.8 | 17.8 |
|  | *Oscillospiraceae* | 15.6 | 13.8 | 14.7 |
|  | *Prevotellaceae* | 11.8 | 3.3 | 7.6 |
|  | *Rikenellaceae* | 6.1 | 3.9 | 5.0 |
| Sessile serrated lesion (Lee et al.) | *Lachnospiraceae* | 30.0 | 30.5 | 30.3 |
|  | *Oscillospiraceae* | 21.5 | 15.2 | 18.3 |
|  | *Bacteroidaceae* | 15.2 | 14.2 | 14.7 |
|  | *Bifidobacteriaceae* | 2.1 | 11.9 | 7.0 |
|  | *Rikenellaceae* | 7.0 | 3.7 | 5.4 |

### Microbial taxonomic differences between FIT NN and FIT SSL participants

Our cohort FIT samples retained 506 of 1,176 species after >10% prevalence filtering (Table 3). Both groups were dominated by the same two phyla, *Firmicutes* and *Bacteroidota*, and at the genus level the same filter retained 312 genera, of which 281 were shared between groups (Figure 4a, Table 3). At the community level, the two groups were indistinguishable. Alpha diversity was equivalent across the Shannon, Simpson and observed-richness indices (p = 0.84, 0.75, 0.21), ordination of Bray-Curtis and Aitchison distance produced overlapping groups, and PERMANOVA attributed less than 6% of variance (R2=0.059, 0.046) to disease status under either metric (Bray-Curtis p = 0.071; Aitchison p = 0.303), with homogeneous within-group dispersion (Figure 4b,c). SSL therefore exhibits a similar overall faecal community to NN, suggesting that any SSL signature in this cohort is likely confined to a small number of individual features rather than a broad shift in composition.

**Figure 4.**
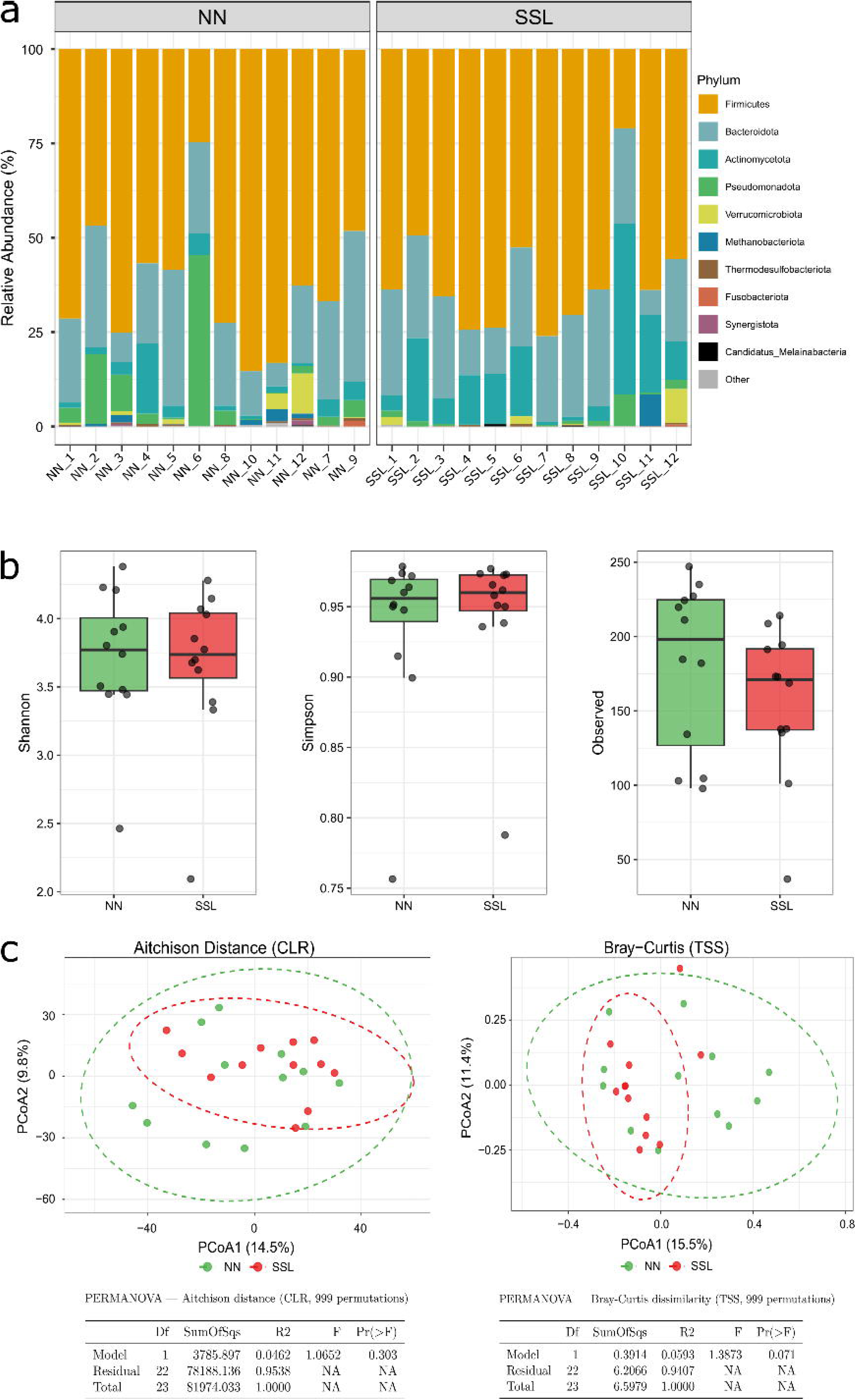
Faecal community structure does not differ between donors with and without a sessile serrated lesion. **a)** Phylum-level relative abundance for each sample, faceted by group. **b)** Shannon diversity, Simpson diversity and observed species richness for 12 SSL and 12 neoplasia-free (NN) FIT-collected samples. Wilcoxon rank-sum p-values are displayed above each comparison and no comparison reached significance. **c)** Principal coordinates analysis of Aitchison distance on CLR-transformed profiles (*left*), and Bray-Curtis dissimilarity on TSS-normalised profiles for the 506 species passing the 10% prevalence filter (*right*). Ellipses denote 95% confidence intervals and PERMANOVA results (adonis2, 999 permutations) are inset. SSL, sessile serrated lesion; NN, neoplasia-free.

**Table 3.** Taxa detected and retained by group. Detected counts are taxa with non-zero relative abundance; retained counts are those passing the >10% prevalence filter used in all downstream analyses. Shared denotes taxa present in both groups; Total is the count in either group.

| Rank | NN | SSL | Shared | Total (either) |
|---|---|---|---|---|
| Detected (before prevalence filter) |  |  |  |  |
| Species | 984 | 757 | 565 | 1,176 |
| Genus | 645 | 487 | 368 | 764 |
| Family | 252 | 180 | 159 | 273 |
| Retained (greater than 10% prevalence filter) |  |  |  |  |
| Species | 491 | 469 | 454 | 506 |
| Genus | 305 | 288 | 281 | 312 |
| Family | 127 | 120 | 118 | 129 |

### Exploratory count-based models suggest directional taxonomic trends in SSL

Applying ALDEx2, we identified none of the 506 retained species were significantly differentially abundant following FDR correction. However, we investigated directional trends across observed taxonomic shifts by applying DESeq2 and edgeR; these tools identified 122 and 65 taxa respectively with generally consistent fold-change estimates (Spearman r = 0.667). Forty-nine were called by both, 36 lower and 13 higher in SSL (Table 4; Figure 5). Taxa lower in SSL were predominantly fermentative commensals, including *Segatella copri* (formerly Prevotella copri), *Parabacteroides goldsteinii* and *Christensenella minuta*, while *Bifidobacterium catenulatum*, *Sellimonas intestinalis* and *Enterocloster clostridioformis* led those higher in SSL. The largest fold changes (log2FC > 20) belong to taxa near-absent in one group and reflect the pseudo-count rather than measured abundance. While the null ALDEx2 result suggests these 49 taxa are not individually differentially abundant, the observed directional changes can inform functional trends across disparate taxa.

**Figure 5.**
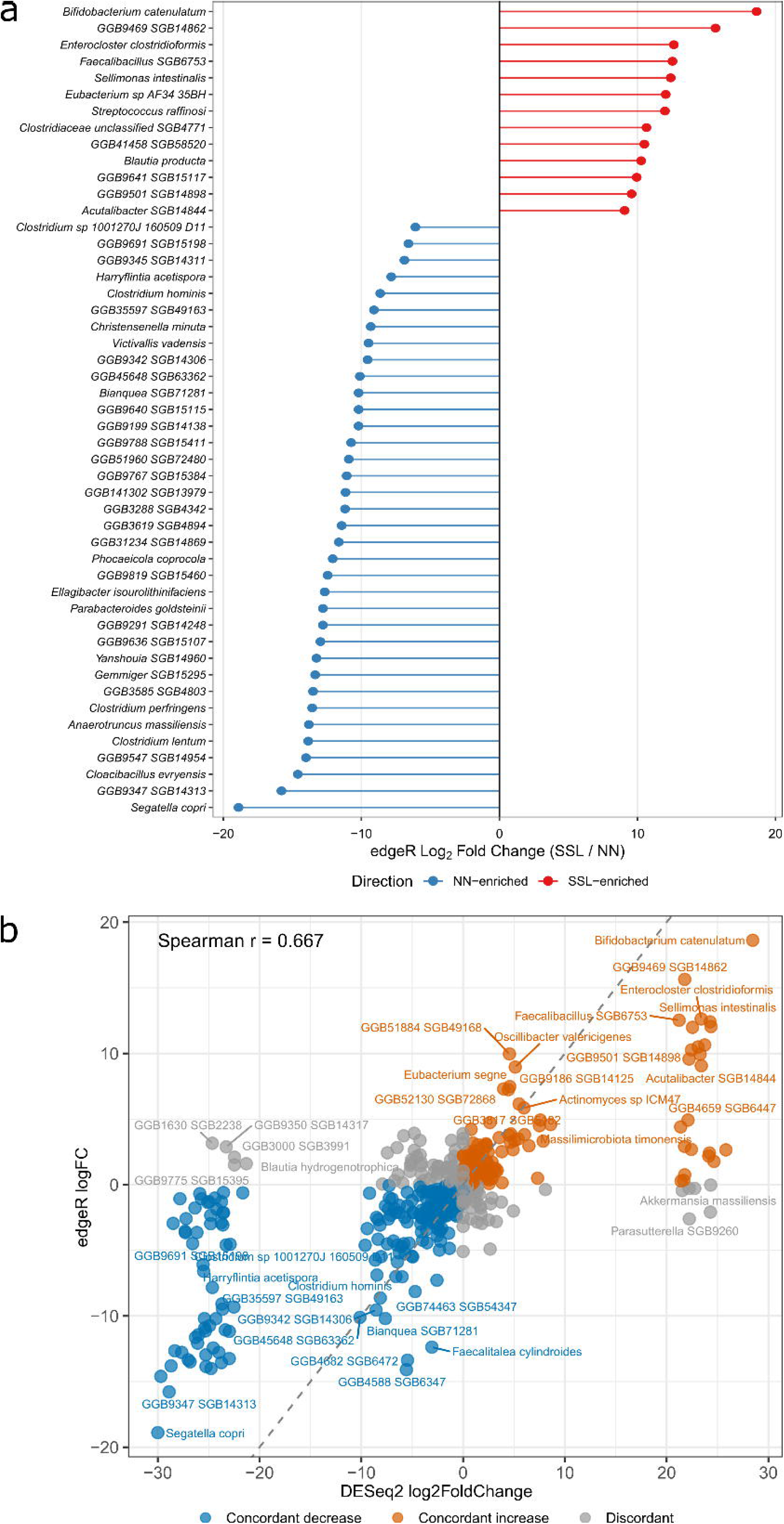
Species called by both DESeq2 and EdgeR. **a)** Lollipop plot of edgeR log2 fold change (SSL/NN) for the 49 consensus taxa. Each species-level genome bin is a stem from zero to its estimate, coloured red for SSL-enriched (positive) and blue for NN-enriched (negative), ordered strongest SSL to strongest NN. SSL enrichment is led by *Bifidobacterium catenulatum* (near +19) and NN enrichment by *Segatella copri* (near −19). **b)** DESeq2 log2 fold change (x-axis) against edgeR log2 fold change (y-axis) for all 506 taxa tested, one point per taxon. Colour denotes cross-method agreement: concordant decrease (blue), concordant increase (orange) and discordant (grey). The dashed diagonal marks exact agreement. The two methods correlate moderately (Spearman r = 0.667), with concordant taxa lying along the diagonal and discordant taxa clustering near the origin where effects are small.

**Table 4.** Consensus directional trending taxa. Only taxa significant under both DESeq2 and edgeR are shown. Top taxa by effect size in each direction are shown; 49 were called by both methods (36 lower, 13 higher in SSL). log2FC from DESeq2, SSL relative to NN.

| Species | log2FC | Direction |
|---|---|---|
| *Bifidobacterium catenulatum* | +28.5 | Enriched in SSL |
| *Eubacterium sp. AF34-35BH* | +24.4 | Enriched in SSL |
| *Sellimonas intestinalis* | +24.3 | Enriched in SSL |
| *Enterocloster clostridioformis* | +23.4 | Enriched in SSL |
| *Streptococcus raffinosi* | +22.5 | Enriched in SSL |
| *Blautia producta* | +22.4 | Enriched in SSL |
| *Segatella copri* | -30.0 | Depleted in SSL |
| *Cloacibacillus evryensis* | -29.7 | Depleted in SSL |
| *Anaerotruncus massiliensis* | -28.7 | Depleted in SSL |
| *Parabacteroides goldsteinii* | -27.7 | Depleted in SSL |
| *Christensenella minuta* | -22.5 | Depleted in SSL |

### SSL microbiomes carry reduced functional richness

HUMAnN3 profiling of the same 24 metagenomes retained 391 MetaCyc pathways and 3,541 KEGG orthologs (KOs) after filtering (Table S4). Although taxonomic community structure did not differ, functional richness was lower in SSL, with fewer KOs (median 2,140 versus 3,160; p = 0.007), fewer pathways (304 versus 374; p = 0.021) and lower KO-level Shannon diversity (6.72 versus 6.94; p = 0.014) (Table 5; Figure 6). Pielou evenness was unchanged at both levels (pathway p = 0.932; KO p = 1.000), so the reduction reflects lost functional breadth rather than dominance by a few abundant orthologs. KO richness tracked *Enterobacteriaceae* abundance closely (Spearman rho = 0.92), and orthologs carried only by *Enterobacteriaceae* added a median of 521 KOs to each NN sample and none to SSL samples.

**Figure 6.**
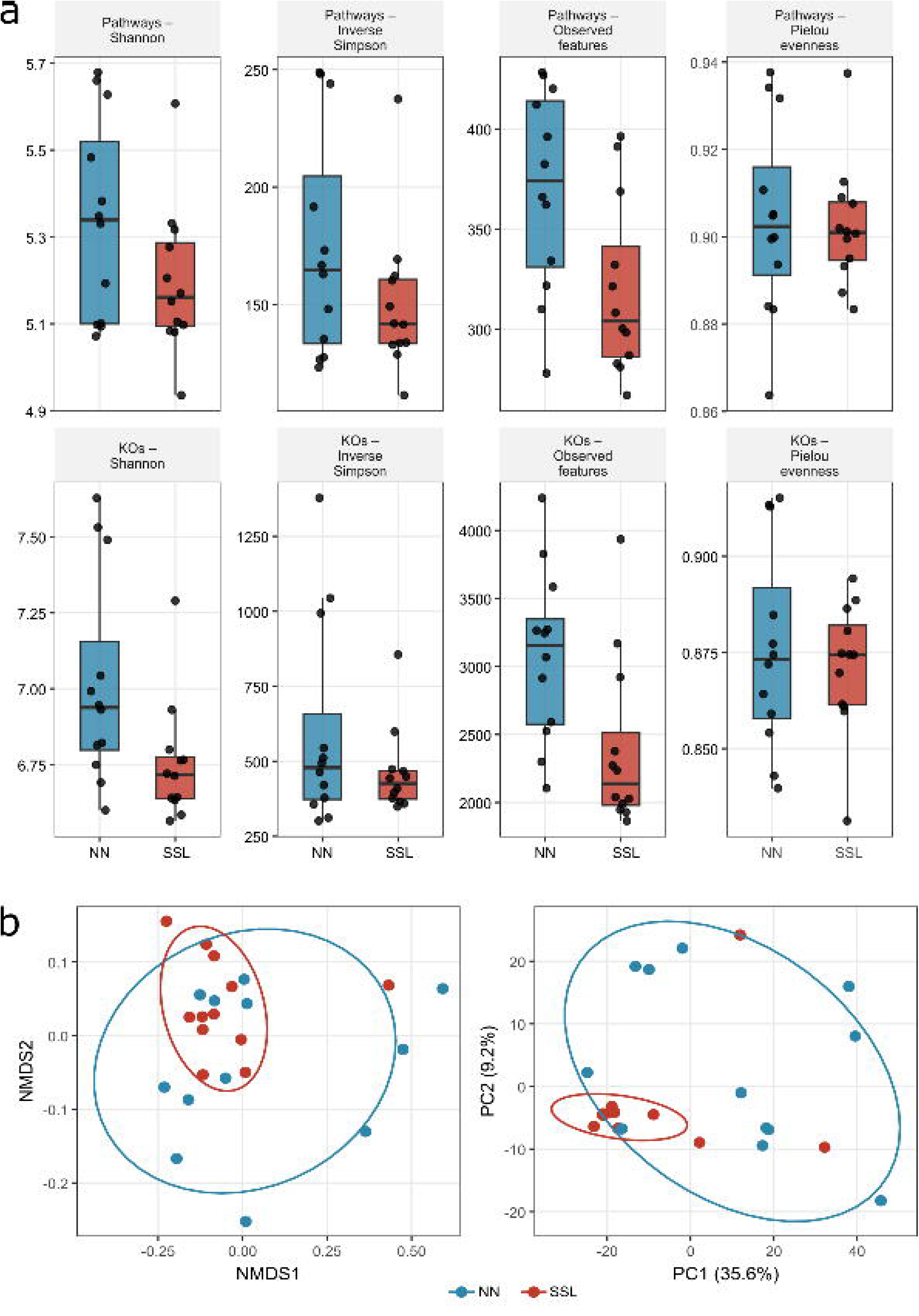
Microbial functional diversity in SSL and NN. **a)** Boxplots of functional alpha diversity indices for MetaCyc pathways (upper panels) and KEGG orthologs (lower panels) across 12 SSL and 12 NN FIT-collected samples. Metrics shown include observed feature count (richness), Shannon diversity, inverse Simpson diversity, and Pielou evenness where applicable. **b)** Functional beta diversity of HUMAnN3-derived pathway profiles in SSL versus neoplasia-free controls showing: non-metric multidimensional scaling (NMDS) ordination of Bray-Curtis dissimilarity computed on pathway CPM profiles (*left*), Principal components analysis (PCA) of CLR-transformed pathway RPK profiles by Aitchison distance (*right*).

**Table 5.** Functional alpha diversity for MetaCyc pathways and KEGG orthologs. Group medians and Wilcoxon rank-sum p-values. Observed richness and KO Shannon diversity are significantly lower in SSL; evenness is unchanged. Abbreviations: KO, KEGG ortholog.

| Layer | Metric | NN | SSL | p |
|---|---|---|---|---|
| Pathways | Shannon | 5.34 | 5.16 | 0.14 |
| Pathways | Inverse Simpson | 165 | 142 | 0.24 |
| Pathways | Observed features | 374 | 304 | 0.02 |
| Pathways | Pielou evenness | 0.900 | 0.901 | 0.93 |
| KOs | Shannon | 6.94 | 6.72 | 0.01 |
| KOs | Inverse Simpson | 479 | 426 | 0.48 |
| KOs | Observed features | 3,160 | 2,140 | 0.01 |
| KOs | Pielou evenness | 0.870 | 0.874 | 1.00 |

### Functional differences concentrate in *Enterobacteriaceae* and *Bifidobacterium* genes

To locate the functional changes, pathways and orthologs were tested with three compositionally aware tools (LinDA, MaAsLin3 and ANCOM-BC2), retaining features supported by two or more methods (76 pathways and 984 KOs; Table 6, Figure 7 a,b,c). Most consensus features were lower in SSL, but stratified profiles showed that this depletion was largely *Enterobacteriaceae* gene content. For 797 of the 830 KOs lower in SSL, *Enterobacteriaceae* carried at least half of the group difference, and *Escherichia coli* alone carried at least half for 587 KOs (Figure S1). Two NN samples contained *E. coli* at 18.3% and 43.3% relative abundance and account for much of this signal. Pathways for ornithine degradation, the enterobacterial common antigen, ADP-heptose and the TCA cycle, and sulphur-related orthologs such as alkanesulfonate monooxygenase, were each dominated by *E. coli* or *Klebsiella* (Figure S2 a,b). Polyamine biosynthesis II was lower in SSL without attribution to any single species. Among orthologs, 130 met the breadth criteria, and 20 of these were called by all three methods. Eighteen of the 20 orthologs were higher in SSL and were carried by several *Bifidobacterium* species, including sucrose phosphorylase (K00690), a raffinose, stachyose and melibiose transporter (K10117) and cobalamin-independent methionine synthase (K00549) (Figure S2 c,d). Sulphate adenylyltransferase (K00957), a core sulphate-activation gene, was carried by many taxa and did not differ between groups (Figure S2).

**Figure 7.**
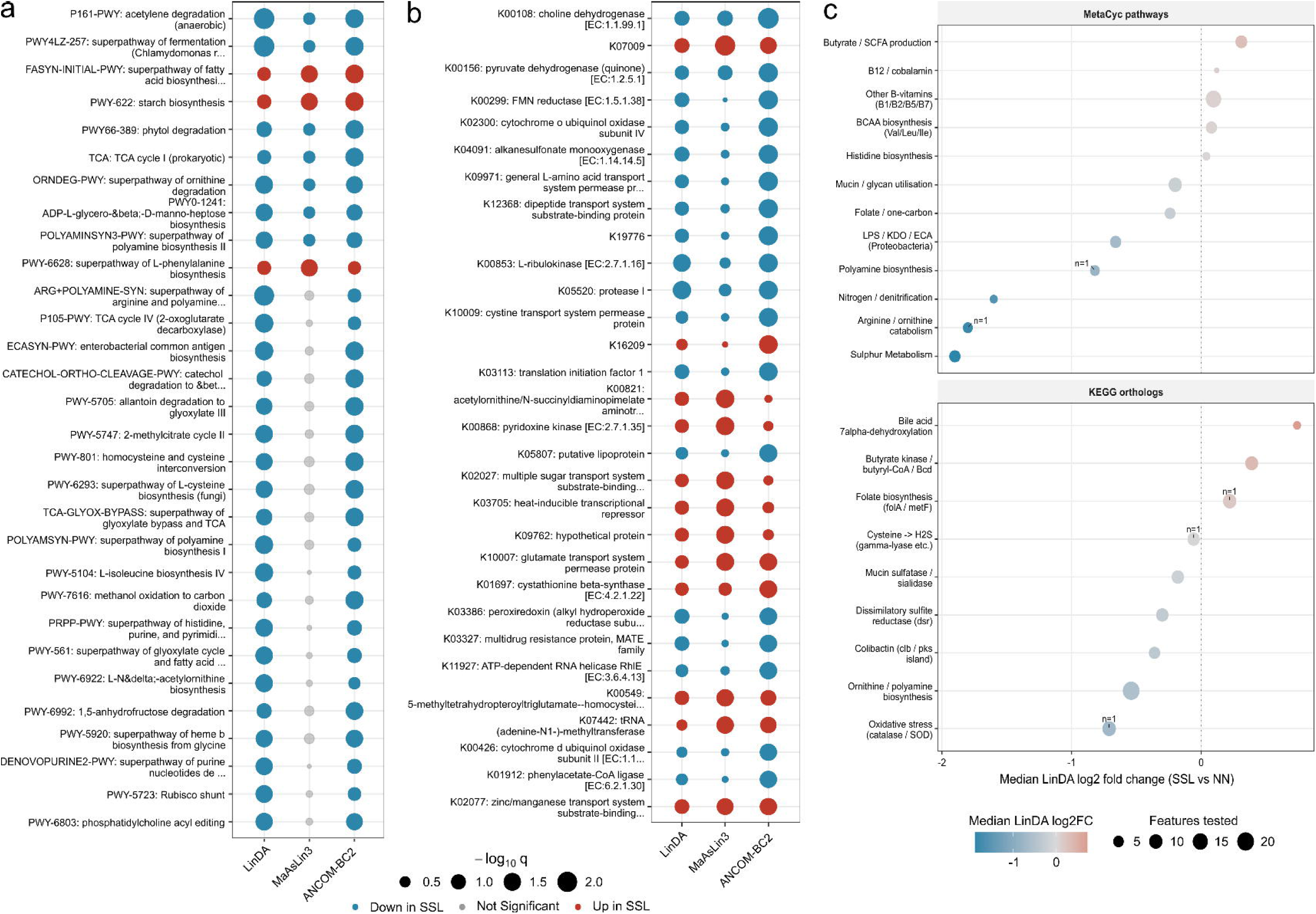
Microbial functional pathway enrichment in SSL and NN. **a)** Dot plot of the top 30 of 166 MetaCyc pathways called by at least one method, with the three methods (LinDA, MaAsLin3 and ANCOM-BC2) on the x-axis and pathways on the y-axis. Dot area scales with significance (−log₁₀ q) and colour denotes the call: blue for down in SSL, red for up in SSL, grey for not significant. **b)** Top 30 of 1,768 KEGG orthologs, ranked by calls across the three methods. **c)** Summary of effect direction by functional category, split into MetaCyc pathways (upper) and KEGG orthologs (lower). The x-axis shows the median LinDA log₂ fold change (SSL versus NN), dot area scales with the number of features tested and colour follows the same fold-change scale (red positive, blue negative). The 166 pathways and 1,768 orthologs shown here are those called by at least one of the three methods, a subset of the 391 pathways and 3,541 orthologs retained after filtering. Categories resting on a single feature are marked n=1. Species attribution of the consensus features is shown in Figure S1.

**Table 6.** Functional consensus features supported by two or more tools (LinDA, MaAsLin3 and ANCOM-BC2). Effect sizes are LinDA log2 fold changes. Main contributors give the largest species and family share of the NN to SSL difference in the HUMAnN3 stratified output; Effective species is the number of equally contributing species that would produce the observed spread of per-species contributions (inverse Simpson index). Pathways without species attribution were reported by HUMAnN3 only at community level.

| Feature | log2FC | Methods | Direction | Main Contributor | Effective Species |
|---|---|---|---|---|---|
| Ornithine degradation (ORNDEG-PWY) | -2.83 | 3 | Down in SSL | *E. coli* 79%; *Enterobacteriaceae* 96% | 1.4 |
| Polyamine biosynthesis II (POLYAMINSYN3-PWY) | -2.02 | 3 | Down in SSL | Not assigned to a species (100% unclassified) | n/a |
| Enterobacterial common antigen (ECASYN-PWY) | -2.21 | 2 | Down in SSL | *E. coli* 65%; *Enterobacteriaceae* 91% | 1.8 |
| ADP-heptose / LPS precursor (PWY0-1241) | -1.43 | 3 | Down in SSL | *E. coli* 57%; *Enterobacteriaceae* 76% | 2.8 |
| TCA: TCA cycle I (prokaryotic) | -2.37 | 3 | Down in SSL | *E. coli* 76%; *Enterobacteriaceae* 93% | 1.4 |
| Branched-chain amino acid transporter ATPase (K01996) | -2.33 | 3 | Down in SSL | *E. coli* 35%; *Enterobacteriaceae* 67% | 4.8 |
| Sucrose phosphorylase (K00690) | +2.50 | 3 | Up in SSL | *B. longum* 40%; *Bifidobacteriaceae* 91% | 3.3 |
| Cobalamin-independent methionine synthase (K00549) | +2.19 | 3 | Up in SSL | *B. longum* 28%; *Bifidobacteriaceae* 77% | 5.0 |
| Raffinose/stachyose/melibiose transporter (K10117) | +1.92 | 3 | Up in SSL | *B. longum* 38%; *Bifidobacteriaceae* 63% | 4.2 |

## DISCUSSION

This study asked whether FIT-collected faeces is suitable for shotgun metagenomic profiling and, if so, whether it can reveal taxonomic or functional differences associated with SSL. In the paired reanalysis of Byrd et al. (13), donor identity explained roughly four times more compositional variance than collection method, although FIT samples consistently carried more *Bacteroidota* than matched fresh samples. In the two cross-cohort comparisons (22,23), alpha diversity did not differ, no species were differentially abundant, and collection method explained at most 7.6% of variance. Within our FIT cohort, SSL and NN samples did not differ in community structure, and no species was differentially abundant under the primary test. Functional richness was, however, lower in SSL. About two thirds of this gap reflected *Enterobacteriaceae* gene content in NN samples and genes carried by several *Bifidobacterium* species were more abundant in SSL.

Reanalysing the paired samples of Byrd et al. (13) allowed the collection-method effect to be measured within each person’s gut microbiome using our pipeline. Donor identity explained 63.5% of Bray-Curtis variance and collection method 16.7%, compared with about 71% and 9% in the original study. A particular concern was that collection method may cause some taxa to not be detected at all. We did identify taxa detected by only one method, however these were all detected at very low abundance in the paired samples, and these ‘absent taxa’ were not unique to one collection method. This likely represents truly lowly abundant species sitting at the threshold for detection and being missed by chance randomly in one method.

The method effect was nonetheless directional; every FIT sample carried more *Bacteroidota* and less *Firmicutes* than its freshly collected pair. The fresh samples carried unusually little *Bacteroidota* (median about 3%), and our FIT samples showed no *Bacteroidota* excess relative to fresh faeces from other cohorts, so part of this shift may lie in the reference arm rather than in FIT. The small compositional shifts in highly abundant taxa we do observe between FIT and fresh are consistent and should not bias comparisons made between FIT samples.

Microbiome profiling is notoriously sensitive to batch effects and comparing samples across studies is difficult. Even so, when our FIT samples were compared with fresh faeces from Piccinno et al. (22) and Lee et al. (23), alpha diversity remained unchanged, no species were differentially abundant, and the two methods produced consistent microbial profiles.

Collection method was significant under PERMANOVA in both comparisons but explained only 6.3% to 7.6% of variance, with no difference in dispersion. Because this figure also absorbs study of origin, it is an upper bound on the FIT effect, and it sits below the 18% study-of-origin variance Pekel et al. (28) found across 6,779 samples. The NN and SSL comparisons behaved alike, so FIT collection does not act selectively on samples from donors with a serrated lesion, and comparisons made within our own FIT cohort should not carry a collection-method bias. FIT-collected samples are therefore suitable for metagenomic comparisons made within FIT-collected cohorts.

Within our own FIT cohort, comparison between SSL and NN samples showed no significant taxonomic differences in alpha diversity, community ordination or ALDEx2 testing. The modest size of this taxonomic signal matches what the literature predicts for an early lesion (4,29,30). In the largest meta-analysis to date, Pekel et al. (28) found adenoma-associated enrichments substantially weaker than the carcinoma signature, with adenoma classifier accuracy falling to an area under the curve of 0.48 to 0.79, so the absence of significant taxonomic differences in a cohort of 24 is what that work would predict for a pre-invasive lesion. The more permissive DESeq2 and edgeR models agreed on 49 species, most of them lower in SSL, and without ALDEx2 support we treat these as directional trends for a larger cohort to test rather than as SSL-associated taxa. *Fusobacterium nucleatum*, among the most strongly documented CRC organisms (31–33), was not among the exploratory SSL taxa, consistent with studies finding no SSL specific enrichment (11,12). The anti-inflammatory commensals *Parabacteroides goldsteinii* and *Christensenella minuta* (34–36) trended lower in SSL, the direction expected of protective bacteria in early carcinogenesis, although neither reached significance under ALDEx2.

The functional results are more informative. SSL samples carried fewer pathways and KOs with unchanged evenness, indicating reduced functional breadth rather than dominance by a few pathways, and the reduced gene richness commonly reported along the conventional adenoma-carcinoma sequence (31,37) appears to extend to the serrated pathway at the level of gene potential. Consensus features lower in SSL included polyamine biosynthesis and a broad set of *Enterobacteriaceae* carried genes for ornithine catabolism, lipopolysaccharide precursors and sulphur metabolism. In the stratified output these losses were attributed to *Escherichia coli* and other *Enterobacteriaceae* rather than to any of the 49 exploratory taxa, so the functional signal is not a restatement of the taxonomic trends. Much of this *Enterobacteriaceae* signal was traced to two NN samples with high *Escherichia coli* carriage, although *Enterobacteriaceae* were also detected in more NN than SSL samples (9 of 12 versus 4 of 12). Genes higher in SSL were instead carried by several *Bifidobacterium* species and centred on oligosaccharide uptake and sucrose metabolism.

Healthy microbial communities typically show functional redundancy, and taxonomic composition can vary widely between individuals while functional capacities are kept by different taxa performing the same roles (38,39). Tian et al. (40) showed that taxa which share gene families form conserved functional networks, so a change in community membership should alter only the functions that few members carry. Our data fits this pattern. Functions carried by many taxa, such as sulphate activation (K00957), did not differ between groups whereas those that did differ were concentrated in single families, with *Enterobacteriaceae* genes lower and *Bifidobacterium* genes higher in SSL. Moya and Ferrer (41) define dysbiosis as a loss of functional capacity beyond what redundancy can buffer.

Because the larger share of the difference is a family whose expansion is itself commonly treated as a marker of dysbiosis (42) and which was more common in NN than SSL samples, we describe the SSL profile as reduced functional breadth rather than early dysbiosis. A cross-sectional design cannot show whether this difference precedes or follows lesion development.

Several functional trends also differ from a late-stage CRC signature. Dissimilatory sulphate reduction genes, the route to hydrogen sulphide that is elevated in CRC (43–45), did not differ between groups, and the sulphur-related orthologs lower in SSL encode sulphur metabolism and cystine uptake by *Enterobacteriaceae*. Superpathway of polyamine biosynthesis II was lower in SSL, opposite to the upregulation reported in colorectal tumours (46). The butyrate producer *Faecalibacterium prausnitzii* was nominally more abundant in SSL (median 2.0% versus 1.5%), and six of seven butyrate and short-chain fatty acid pathways trended higher. Although none reached significance, this is the reverse of their depletion in CRC (47). These trends appear to support a potential early state that precedes the functional changes of carcinoma, although greater statistical power is required to reach significance.

To our knowledge, this is the first study to use FIT-collected samples to profile the taxonomic and functional metagenome of SSL, a lesion under-represented relative to conventional adenomas (10,12). The functional differences suggest that SSL leaves a detectable faecal signature, and the clinical gap it represents is worth addressing. FIT detects sessile serrated adenomas or polyps with a sensitivity at or below 12.3% across haemoglobin cut-offs (5,48), yet the serrated route contributes substantially to sporadic and interval cancers (29,30,49). A FIT-based metagenomic assay would use samples already collected at population scale, with no change to the screening workflow. Because gene content revealed differences that taxonomic tests did not, such a classifier should prioritise multi-feature functional axes carried by several taxa over any single taxon or any function supplied by one organism.

Linking functional differences to the taxa that carry them also suggests a practical route to a diagnostic. Shotgun sequencing of every screening sample is unlikely to be affordable at population scale, but a functional signature traceable to a defined panel of carrier taxa could be measured directly in the FIT sample with targeted assays such as quantitative PCR. Quantifying *Fusobacterium nucleatum* in FIT samples has already improved CRC detection beyond faecal haemoglobin alone (50). In this cohort, functions higher in SSL traced to several *Bifidobacterium* species and those lower in SSL mainly to *Enterobacteriaceae*. Neither group is yet a candidate marker, because both vary with diet and medication and the *Enterobacteriaceae* signal rested heavily on two control samples.

We acknowledge several limitations that may affect interpretation of the results. In the cross-cohort comparisons, collection method cannot be separated from study of origin, and no formal equivalence test was performed. The cohort was small (n = 12 per group) with limited statistical power. The primary taxonomic test returned no significant species, so we consider trends from the more permissive models as exploratory. The two NN samples dominated by *Escherichia coli* drive the functional pathways supported by all three differential abundance methods, so the depletion of *Enterobacteriaceae*-carried functions may be unique to this cohort. The SSL group was predominantly female and had a higher median BMI than the NN group, and five participants with an SSL also carried a synchronous advanced adenoma, so part of the signal may reflect these differences rather than the SSL itself. Metagenomic sequencing infers functional potential rather than activity (51). Gene content remains informative, however, as unutilised genes confer a negative selection pressure, so conserved functions should generally reflect those necessary in that environment. Whether the FIT buffer preserves RNA well enough for metatranscriptomics is unknown and would need direct testing. For SSL, the next step is to confirm a functional signature in larger multi-centre screening cohorts and test whether measuring its carrier taxa at the time of FIT improves detection of the lesions that FIT alone misses.

## Conclusions

FIT stabilised faeces is a suitable sample source for shotgun metagenomic biomarker discovery, with collection method contributing far less variance than differences between individuals. Within FIT samples, SSL and NN groups did not differ in community structure or in any individual species, but SSL samples carried fewer functional pathways and orthologs, including lower polyamine biosynthesis capacity and functional signatures driven by several *Bifidobacterium* species. A multi-feature functional and taxonomic signature, confirmed in larger cohorts, offers a route to detecting SSLs from samples that screening programs already collect.

## Funding Statement

This work is supported by funding from the National Health and Medical Research Council (NHMRC) Ideas Grant 2012689, a Flinders Foundation Health Seed Grant, and further funding support from the Flinders Accelerator for Microbiome Exploration. MJR is supported by NHMRC Ideas Grant 2036844. JMW is supported by a Cancer Council SA Research Fellowship. Eiken Chemical Co. Ltd. for provision of the FIT devices and OC-Sensor faecal testing devices

## Acknowledgements

We would like to thank the participants for providing their samples for the research study. The authors acknowledge the provision of computing and data resources provided by the Australian BioCommons Leadership Share (ABLeS) program. This program is co-funded by Bioplatforms Australia (enabled by NCRIS), the National Computational Infrastructure and Pawsey Supercomputing Research Centre (52).

## Declaration of Interest Statement

E.L.S. reports receiving institutional funding and consumables from Eiken Chemical Company for investigator led studies on colorectal cancer screening research. All other authors report no conflicts of interest.

## Data Availability Statement

All raw sequencing data generated in this study have been submitted to the NCBI Sequence Read Archive (ncbi.nlm.nih.gov/sra) under NCBI BioProject accession PRJNA1522997.

Human sequences have been removed prior to deposition. All bioinformatics code used in this study is available at github.com/biobrad/Metagenomic_Landscapes_SSL_Pathway_Scripts and doi.org/10.5281/zenodo.23965408. Supplementary tables are available at doi.org/10.5281/zenodo.23966270.

## Author Contributions

Conceptualization: GLL, JMW, ELS, MJR; Funding acquisition: MPD, RAE, JMW, ELS; Investigation: BJH, GLL, MJR; Methodology: CC, BJH, GLL, JMW, MJR; Data curation: BJH, RP, MM, GLL, MJR; Formal analysis: BJH; Resources: RAE, MPD, GLL, CC, JMW, ELS; Project administration: JMW, ELS, MJR; Visualization: BJH; Supervision: JMW, ELS, MJR; Writing–Original Draft Preparation: BJH; Writing–Review & Editing: all authors.

## Supplementary Tables

**Table S1:** Definitions of sessile serrated lesions

**Table S2:** Sequencing information

**Table S3:** [paired FIT vs direct samples] taxa detected in one method

**Table S4:** Functional Features and filtering

**Table S5:** Species attribution and breadth criteria for all functional consensus features Table S6: Comparability of FIT and freshly collected faeces metagenomes across two cohorts.

## Supplementary Figures

**Figure S1.**
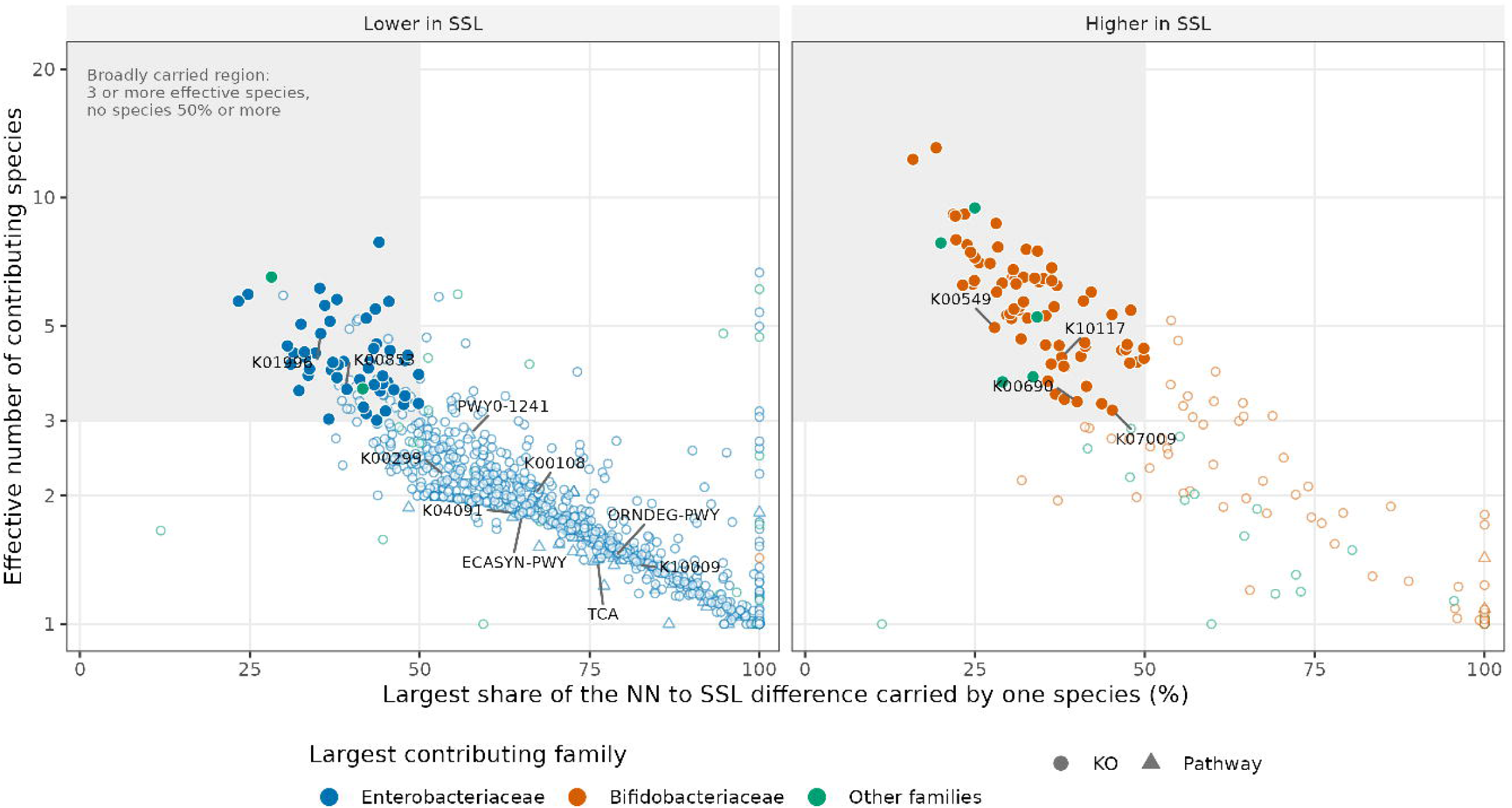
Taxonomic breadth of the functional consensus features. Each point is a MetaCyc pathway (triangle) or KEGG ortholog (circle) called by at least two of LinDA, MaAsLin3 and ANCOM-BC2. The x axis shows the share of the NN to SSL difference carried by *Enterobacteriaceae* in the HUMAnN3 stratified output; the y axis shows the effective number of contributing species. Filled points meet all three criteria for a broadly carried feature (shaded region: three or more effective species). Colour gives direction: blue lower in SSL, red higher in SSL. Features discussed in the text and Table 6 are labelled.

**Figure S2.**
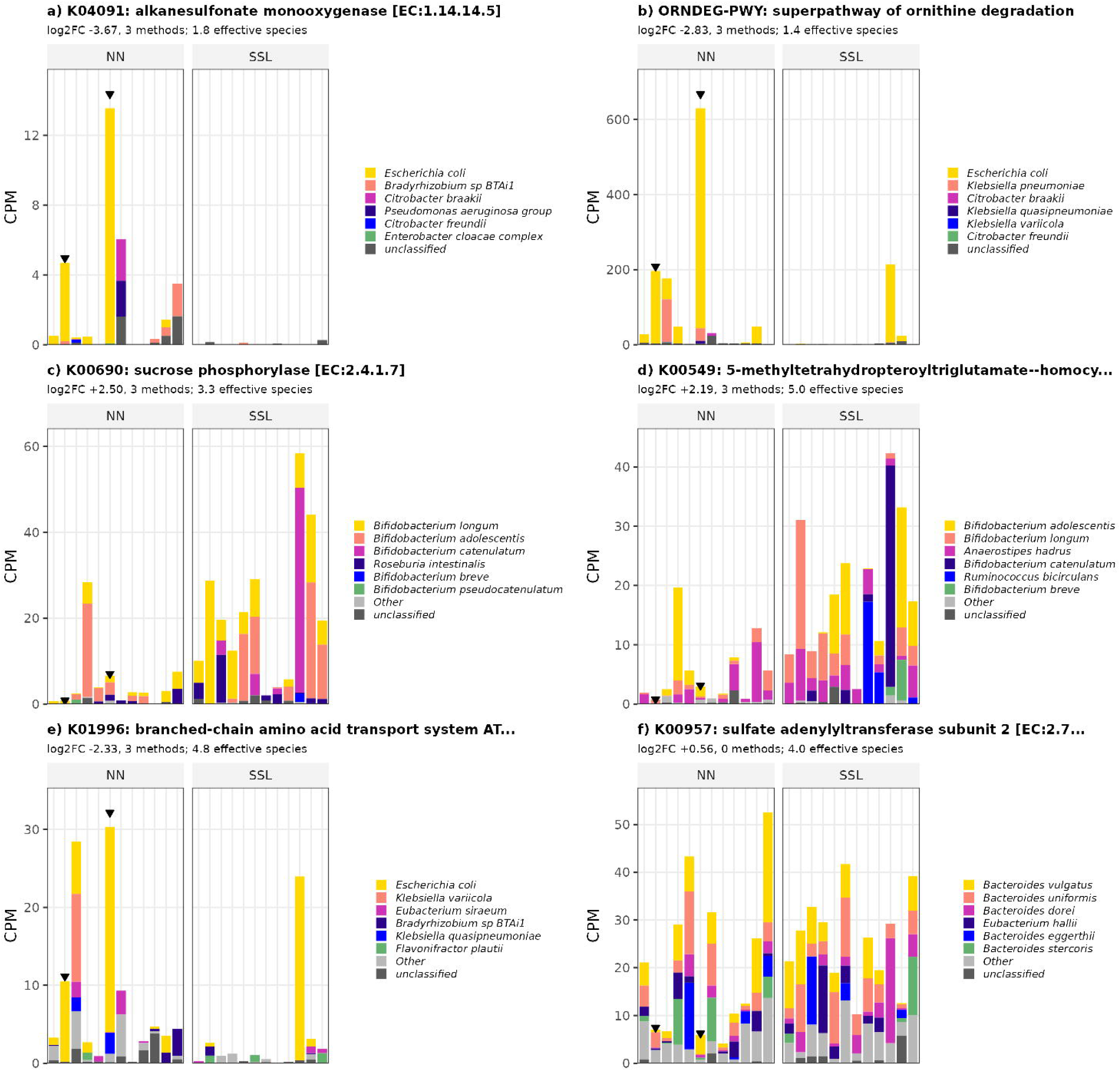
Per-sample contributions of individual species to selected functional features. Each bar is one sample, grouped by NN then SSL; the six largest contributing species are shown and the remainder pooled as Other. Triangles mark the two NN samples with the highest *Escherichia coli* abundance. a) Alkanesulfonate monooxygenase (K04091) and b) ornithine degradation (ORNDEG-PWY), consensus features whose difference is carried by *E. coli* and other *Enterobacteriaceae*. c) Sucrose phosphorylase (K00690) and d) cobalamin-independent methionine synthase (K00549), consensus orthologs higher in SSL and carried by several *Bifidobacterium* and other species. e) Branched-chain amino acid transporter ATPase (K01996), lower in SSL and spread across several *Enterobacteriaceae* and other species. f) Sulphate adenylyltransferase subunit 2 (K00957), carried by many taxa and not different between groups.

